# A gap-free telomere-to-telomere pig reference genome provides insight in centromere evolution and breed-specific selection

**DOI:** 10.1101/2025.10.02.679976

**Authors:** Dong Li, Yulong Wang, Tiantian Yuan, Xiang Li, Yu Quan, Wansheng Liu, Kyle M. Schachtschneider, Lawrence B. Schook, Richard P.M.A. Crooijmans, Martien A.M. Groenen, Martijn F.L. Derks, Taiyong Yu

**Affiliations:** College of Animal Science and Technology, Northwest A&F University; Yangling, Shaanxi, 712100, China; Animal Breeding and Genomics, Wageningen University and Research; Wageningen, 6700 AH, The Netherlands; Department of Animal Science, Center for Reproductive Biology and Health (CRBH), College of Agricultural Sciences, The Pennsylvania State University; University Park, 16802, PA, USA; Sus Clinicals Inc; Chicago, 60612, IL, USA; Department of Animal Sciences, University of Illinois at Urbana-Champaign; Urbana, 61801, IL, USA; Department of Radiology, University of Illinois at Chicago; Chicago, 60612, IL, USA

## Abstract

The domestic pig is a key agricultural species and biomedical model, yet its reference genome has remained incomplete. Using PacBio HiFi, Oxford Nanopore reads, and Hi-C sequencing, we assembled the first telomere-to-telomere (T2T), gap-free pig genome (T2T-Sscrofa). Importantly, this assembly derives from the same individual pig that provided the original reference genome, thereby completing the long-standing foundation of pig genomics. This assembly resolves 274.8 Mb of previously unassembled sequence, including centromeres, segmental duplications, and ribosomal DNA arrays, and identifies 255 new protein-coding genes. Comparative analyses reveal a Robertsonian translocation in the western European wild boar and extensive structural variation across global pig populations. T2T-Sscrofa provides a comprehensive genomic foundation for agricultural, evolutionary, and biomedical research.

## Introduction

The domestic pig (*Sus scrofa* domesticus) is one of the most common mammals worldwide and the primary source of pork, the most consumed meat globally (*1*). Beyond their agricultural importance, pigs are also widely utilized as biomedical models, making their genome central to progress in food security and scientific research. The International Swine Genome Sequencing Consortium (ISGSC) initiated the genome sequencing project from a single female Duroc pig that was raised at the University of Illinois (*2*). The draft pig genome was published in 2012 (*3*), and the current pig reference genome was released in 2017 (*Sscrofa*11.1) (*4*). Hence, this reference traces its origin to 2005 and has been continually improved over the past two decades. The most recent pig reference genome (*Sscrof*a11.1) spans 2.5 gigabases (Gb) across 705 scaffolds and contains 93 gaps. Many of these unresolved regions, particularly centromeres on acrocentric chromosomes, remain inaccessible due to complex structural variation (SV) and satellite repeats, resulting in unfinished or misassembled genomic segments (*5*). Long-read sequence technology such as PacBio circular consensus sequencing (HiFi) and Oxford Nanopore Technologies (ONT) ultra-long sequencing have made it possible to overcome past genome assembly challenges and achieve telomere-to-telomere (T2T) genome assemblies (*6*). Here, we assembled the initial T2T gap-free pig genome (T2T-Sscrofa), the current version covers previously unknown regions, reveals previously unidentified genes, and corrects sequencing errors present in *Sscrofa*11.1.

## Results

### The complete genome of Sus scrofa

T2T-Tabasco was extensively sequences with multiple technologies, including 276 Gb PacBio HiFi, 395 Gb ONT ultra-long, 606 Gb High-throughput Chromosome Conformation Capture (Hi-C) (**Table S1**). T2T-Tabasco includes gapless telomere-to-telomere assemblies for all 18 pig autosomes and chromosome (chr) X, with Hi-C matrix providing independent validation of the assembly structure (**Fig. S1**). PacBio HiFi K-mer analysis estimated the assembly to be highly accurate (QV = 68.8) (**Table S2**) (*7*). Since Tabasco is a female, we used a complete Y chromosome from another Duroc (GCA_023065335.1) as the Y chromosome for the final T2T-Sscrofa. The final genome is 2,688,404,542 bp (**Fig. 1A and Table S3-S4**). Comparison between the T2T-Sscrofa and *Sscrofa*11.1 assemblies revealed high collinearity (**Fig. S2**) (*4*), and a total of 274.8 megabase (Mb) of previously unassembled regions (PURs), ranging in length from 3.8 to 33.3 Mb on different chromosomes (**Fig. 1B and Table S5**). Analysis of the additional sequences in the T2T assembly revealed that PURs was composed of centromeric sequences (137.9 Mb, 50.1%), segmental duplications (SDs) (66.1 Mb, 24.1%), SDs overlapping centromeres (4.6 Mb, 1.7%), other repeat sequences (47.3 Mb, 17.2%), and unclassified sequences (18.9 Mb, 6.9%) (**Fig. 1C**).

**Fig. 1.**
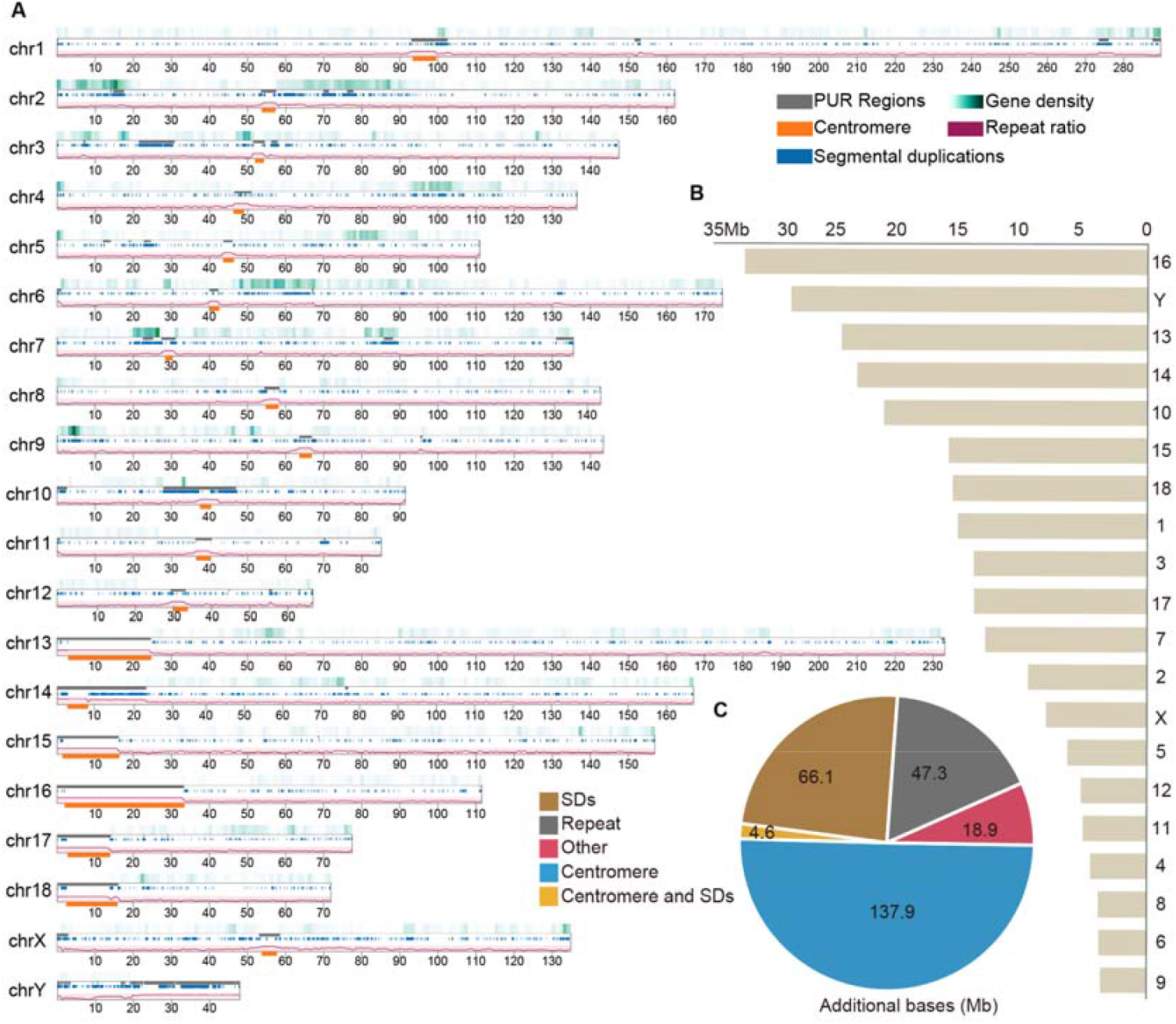
Summary of the telomere-to-telomere pig genome assembly. **A**. Ideogram of T2T-Sscrofa assembly features. For each chromosome (chr), the following tracks are displayed from outer to inner: previously unassembled regions (PURs) in black; gene density gradient from white (low) to green (high); centromeres in orange; repeat content shown as purple curve; segmental duplications (SDs) in blue. Scale bar indicates megabase (Mb). **B**. Additional (nonsyntenic) sequences in the T2T-Sscrofa assembly relative to *Sscrofa*11.1 reference genome, quantified per chromosome in Mb. **C**. Composition of the additional sequences.

### Centromere and rDNA analysis

Chr1-12 and chrX are metacentric/submetacentric chromosomes, whereas chr13-18 are acrocentric (**Fig. 1B**) (*8*). The centromere length of metacentric/submetacentric chromosomes ranges from 2.5 to 6.5 Mb (**Table S6**). Nine centromeric (satellite) repeat sequences were found to be abundant in the pig centromere (**Table S7**), For chr1-4, chr6, chr12, and chrX, the pig centromeric repeats (pigcent) are predominantly composed of pigcent2 (44.2-49.4%), pigcent3 (0.9-15.7%), and pigcent5 (37.1-46.4%). Chr5 and chr7-11 are characterized by pigcent2 (5-29.9%), pigcent5 (5.6-21.9%), pigcent7 (16.6-36.4%), pigcent8 (12.7-30.6%), and pigcent9 (2.3-20.9%) (**Fig. 2A and Table S8**). Interestingly, acrocentric chromosomes in T2T-Sscrofa harbor substantially longer centromeres than metacentric/submetacentric chromosomes, reflecting the accumulation of repetitive sequences, a pattern observed in other mammals (*9*). Acrocentric short arms vary from 1.3 to 2.7 Mb in length, and the centromere ranged from 5.4-31.6 Mb (chr13-chr18) (**Fig. 2B**). The acrocentric centromere repeats are mainly composed of pigcent4 (33.8-44.9%), pigcent5 (14.7-24.3%) and pigcent6 (26.9-49%), distinct from metacentric/submetacentric centromere satellite sequences. The longest centromere spans 31.6 Mb on chr16 (**Fig. 2C**). Pigcent4-6 represents the major component of centromeric regions on the acrocentric chromosomes. This region is characterized by high repetitive content, weak Hi-C interaction signals, stable DNA methylation ratios, sparse distribution of Short Interspersed Nuclear Elements (SINEs) and Long Interspersed Nuclear Elements (LINEs), and an absence of detectable genes. The short arms of the acrocentric chromosomes contain SDs, but no genes or ribosomal DNA (rDNA) sequences were annotated in these regions, different from the rDNA clusters observed in human short arm acrocentric chromosomes (*10*).

**Fig. 2.**
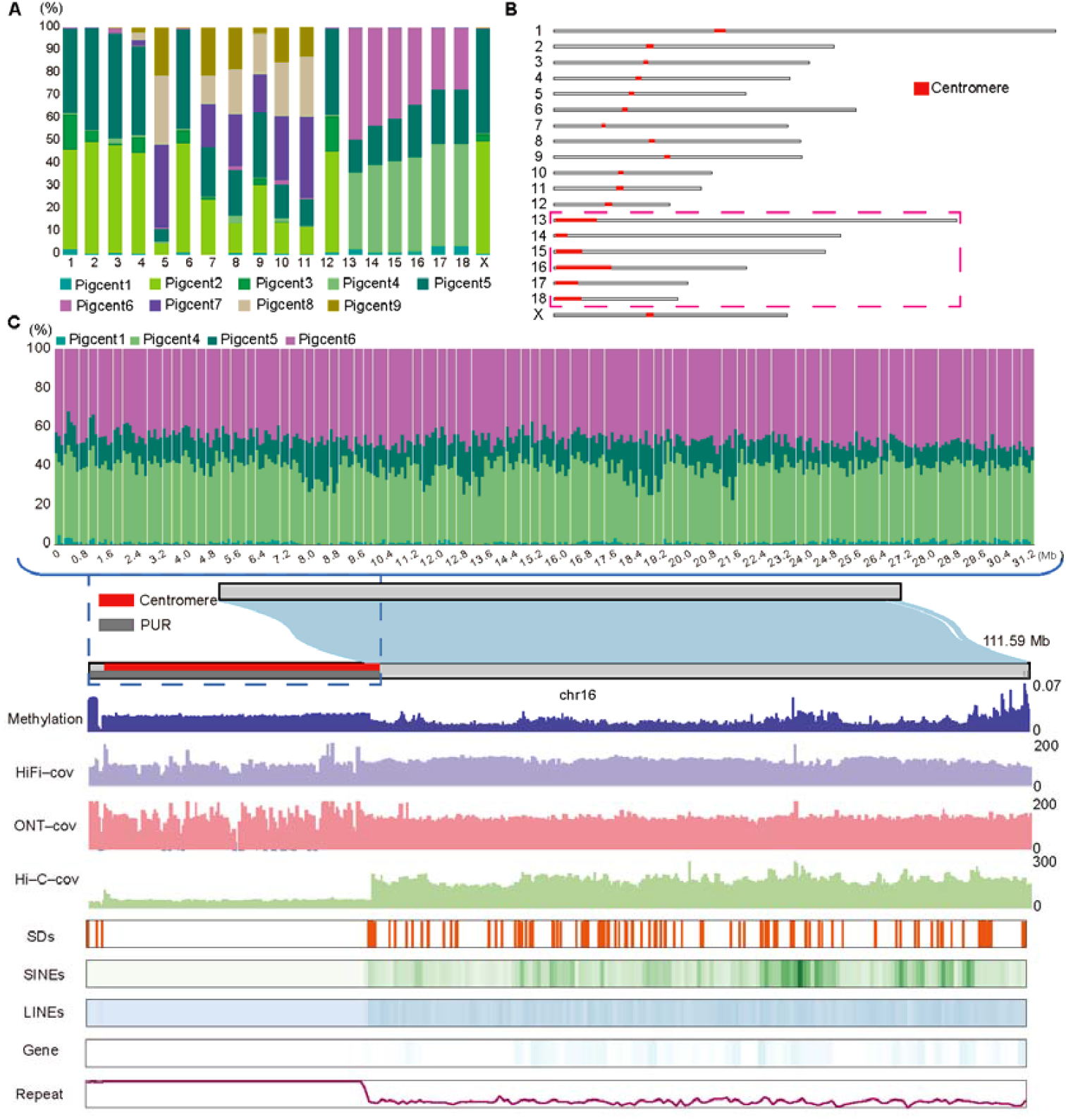
Centromere analysis of T2T-Sscrofa. **A**. Centromeric satellite sequence composition of different chromosomes. **B**. Distribution of centromere sequences on different chromosomes. **C**. Genomic features of chromosome 16. From top to bottom, tracks display: methylation proportion in 10kb windows; sequencing coverage for PacBio HiFi (HiFi-cov), Oxford Nanopore ultralong (ONT-cov), and Hi-C (Hi-C-cov) in 100kb windows; density of short interspersed nuclear elements (SINEs, green) and long interspersed nuclear elements (LINEs, blue) with gradient from white (low) to colored (high); total repeat content (purple curve); and segmental duplications (SDs, red bars). Each track represents the relative abundance or density of genomic features across chromosomal intervals.

rDNA arrays are mainly concentrated on chr8, chr10 and chr14 (59, 20, 148 respectively) (**Table S9**) consistent with findings in other high quality pig genome assemblies (**Fig. S3 and Table S10**). Chr8 exhibits a tandem distribution of 5.8S, 18S, and 28S rDNA repeats. Chr10 displays a bipartite organization, with one region showing the same distribution pattern as chr8, while the other region contains only 18S type repeats. Chr14 is characterized exclusively by 5S repeat sequences.

### Genome annotation analysis

The T2T-Sscrofa genome has a GC content of 42.6%, slightly higher than *Sscrofa*11.1 (41.9%), with 1,112 Mb (41.4%) and 911 Mb (36.4%) of sequence masked, respectively (**Table S11**). Unclassified repeats increased from 25.5% to 29.3%, reflecting improved capture of complex repetitive regions. Interspersed repeats occupy 37.5% of T2T-Sscrofa, including 757,667 retroelements spanning 209 Mb (7.8%), predominantly LINEs (182.1 Mb, 6.8%) of the L1/CIN4 subfamily (6.3%), and SINEs (17.7 Mb, 0.7%). DNA transposons are less abundant, with 39,105 elements covering 10.4 Mb (0.4%).

The public NCBI eukaryotic Genome Annotation Pipeline (Egapx) and transanno (NCBI v106 pig reference annotation) completed the annotation of T2T-Sscrofa by combining *de novo*, homology prediction and RNA-seq evidence (**Table S12**). A total of 21,108 protein coding genes (20,674 in *Sscrofa*11.1) were annotated and 255 are novel protein coding genes (**Table S13**). Furthermore, T2T-Sscrofa assembly placed 887 protein-coding genes that remained unlocalized in the *Sscrofa*11.1 genome particularly on chr1, chr2, chr5, and chr7 (**Fig. S4 and Table S14**).

### Western European wild boar carries a unique chromosome 15-17 Robertsonian fusion

Western European wild boars (EW) carry a unique chromosomal fusion, in which ancestral chr15 and chr17 have joined via a Robertsonian translocation. Despite this structural difference, hybrids between fused and standard karyotypes are observed in the wild and remain fully fertile (*11*). We sequenced a male wild boar from the Netherlands to characterize this fusion event and assembled a chromosome-level EW genome using ONT long reads (122 Gb), Illumina sequencing (32 Gb), and Pore-C (159 Gb). In addition, we assembled an Asian wild boar (AW) and a central EW individual with high quality long reads (**Table S15-17**). The other autosomes of the western EW genome show high collinearity with the 16 autosomes of T2T-Sscrofa (**Fig. 3A**). Chr15 and 17 are fused into a single chromosome in western EW due to a Robertsonian translocation between ancestral chr15 and 17 (**Fig. 3B and Fig. S5)** (*12*), whereas the assembly of AW and central EW did not show the fusion event, as supported by Hi-C contact maps (**Fig. 3B and Fig. S5)**. Furthermore, we aligned 30 high-quality long-read datasets to T2T-Sscrofa (coverage: 40.7-62.5X) (**Table S18**) in which the short-arm regions of chr15 and chr17 showed markedly reduced coverage in western EW, consistent with the short arms being lost during the Robertsonian fusion event (**Fig. 3C and Table S19)**, while coverage across chromosomes 13-18 was comparable to other samples.

**Fig. 3.**
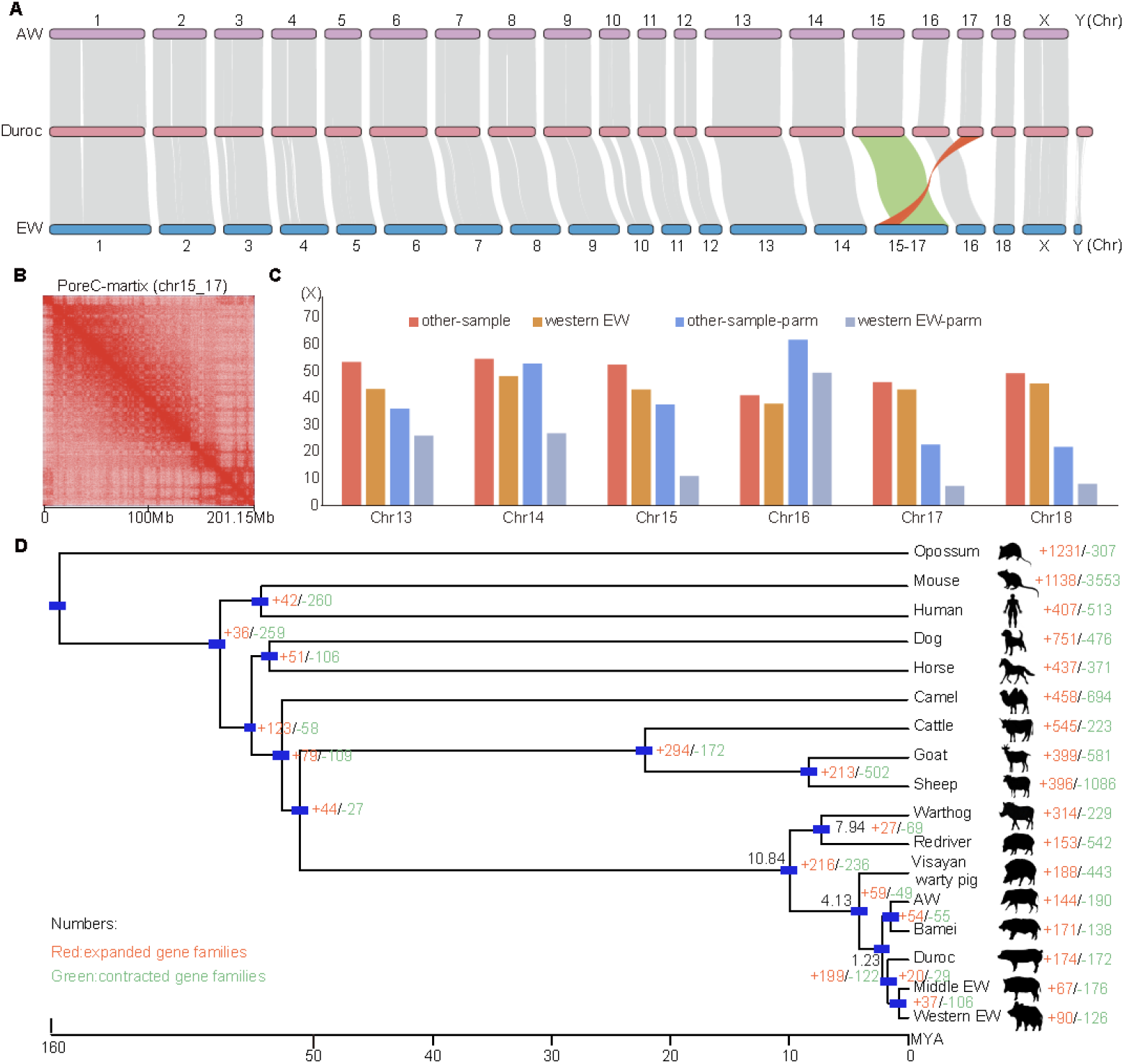
Analysis of western European wild boar. **A**. Collinearity analysis among western European wild boar (EW), Asian wild boar (AW) with T2T-Sscrofa. **B**. The Pore-C chromosome conformation capture (Pore-C) matrix of the chr15_17 sequence of western EW. **C**. Long-read sequencing coverage across chr13-18 and detailed coverage of the short arms of chr13-18, comparing western EW with other samples. **D**. Phylogenetic tree with divergence time estimates. Numbers at nodes indicate divergence times (million years ago, Mya). Gene family dynamics are shown on branches, with numbers indicating expanded (red) and contracted (green) gene families following each speciation event.

### Suidae evolution and domestication

We examined the evolutionary relationships among several *Suidae* species, estimating the divergence time between the African suids *Potamochoerus porcus* (Red river hog) and *Phacochoerus africanus* (Warthog) and other *Sus* species to be 11 million years ago (Mya), while *Sus cebifrons (*Visayan warty pig) and *Sus Sscrofa* diverged about 4.13 Mya. EW and AW diverged about 1.23 Mya (**Fig. 3D and Table S20**), somewhat earlier than previously reported, with domestication later occurring independently in both lineages (*13*). Gene family expansions in domestic pigs versus wild boars showed that expanded genes in domestic pigs were enriched in pathways related to muscle development, cardiac function, ATP metabolism and cellular signaling, supporting enhanced growth, meat quality and reproductive traits targeted during domestication (**Table S21**). Wild boars showed expansions in pathways related to carbohydrate metabolism (oligosaccharide catabolism), ion transport, lipid metabolism, detoxification, and membrane integrity, adaptations that are likely to support survival in diverse wild environments with variable food sources and environmental stressors (**Fig. S6**).

### Structural variation and population genomics across Eurasian pigs

To comprehensively characterize genomic variation across pig populations, we mapped long-read sequences from 45 samples to T2T-Sscrofa, achieving an average mapping rate of 95.9-99.4% (**Table S22**). We detected 46,190-60,295 SVs in European pigs, 81,685-84,419 in Tibetan × Berkshire hybrids, and 65,063-91,913 in Asian pigs (**Fig. 4A**), with precision of 0.89-0.95, recall of 0.84-0.93, and F1 scores of 0.88-0.94 (**Fig. 4B**). After merging, 229,824 SVs were retained, including 126,059 insertions (54.9%, median 288 bp), 101,168 deletions (44.0%, median 290 bp), 1,857 inversions (0.8%, mean 8.3 kb), and 740 duplications (0.3%, mean 13.9 kb) (**Table S23**). Two peaks in the size distribution 201-400 bp (46.2%) and 500 bp-5 kb (15.3%), correspond to the abundant SINEs and LINEs in the pig genome (**Fig. 4C**). We retained 27,939 SVs in PURs (274.8 Mb), including 10,564 insertions (37.82%), 17,155 deletions (61.38%), 66 inversions (0.24%), and 154 duplications (0.55%). Only 2,792 SVs occurred in gene regions, with the majority located in intergenic regions (90%) (**Table S24**).

**Fig. 4.**
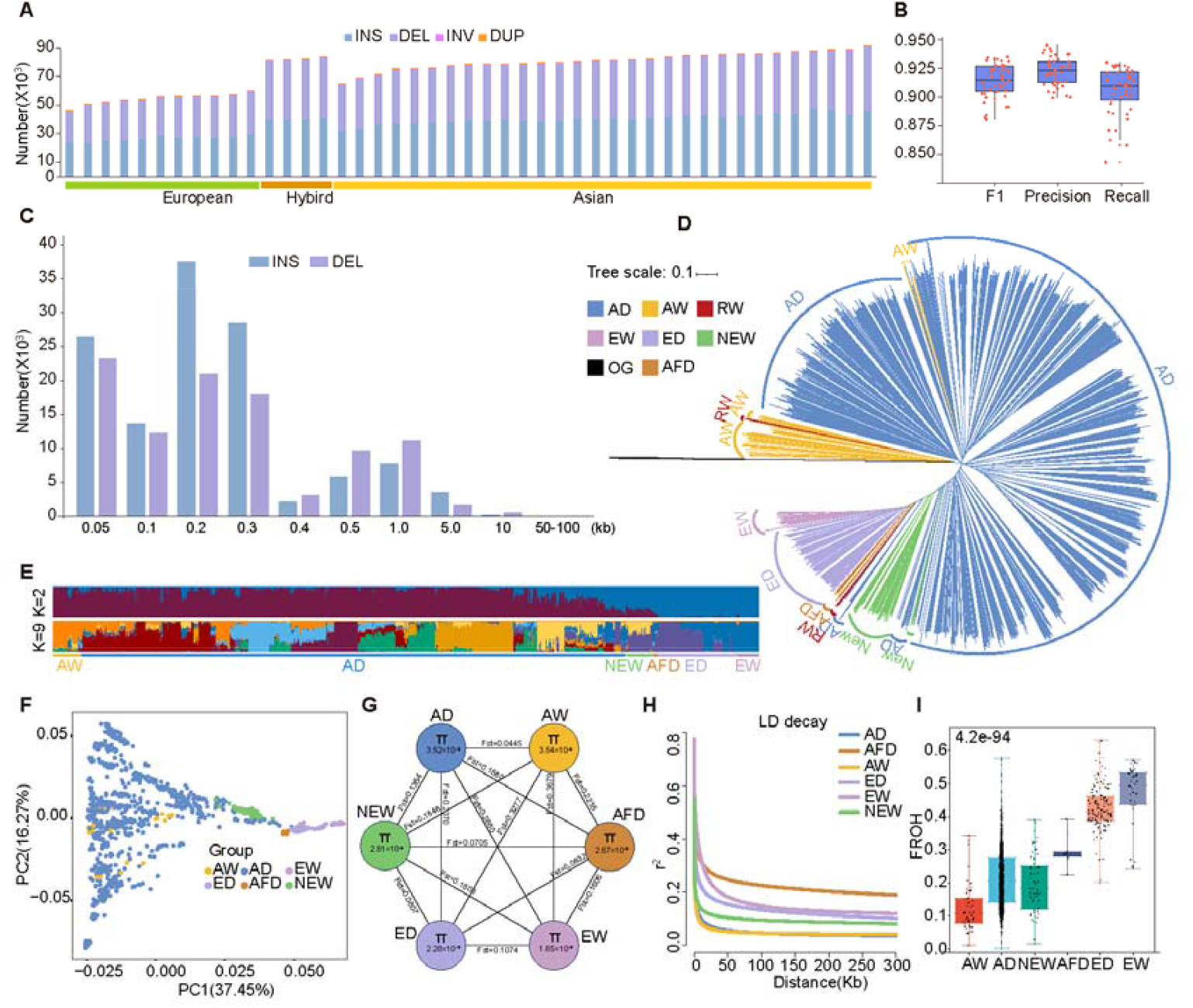
Variant detection and population structure analysis. **A**. Summary statistics of structural variations (SVs). **B**. Performance evaluation of SVs. Box plots show the distribution of F1 score, precision, and recall metrics. **C**. Length distribution of insertion and deletion SVs. **D**. Neighbor-joining tree of wild and domestic pigs based on SNPs. AD: Asian domestic pig; ED: European domestic pig; AW: Asian wild boar; EW: European wild boar; NEW: Newer pig populations formed after the turn of the century; AFD: Africa domestic pig; OG: outgroup. **E**. Population genetic structure of wild and domestic pigs inferred from Admixture (K=2, 9). **F**. Principal Component Analysis (PCA) of pig populations based on SNPs. **G**. Summary of nucleotide diversity (π) and population divergence (Fst) among different pig populations. **H**. Linkage disequilibrium (LD) decay in different pig populations. **I**. Boxplot of the fraction of runs of homozygosity (FROH) among different pig populations.

To further assess genomic variation at the population level, we analyzed 1,303 whole-genome sequencing (WGS) samples (average coverage 96.1%, average depth 18.2X) representing 156 pig populations from diverse geographic regions (**Fig. S7 and Table S25**). We retained 111,457,594 biallelic single nucleotide polymorphisms (SNPs) and 13,249,688 biallelic insertion-deletions (indels), with 1.95% of SNPs and 1.77% of indels located within PURs. On autosomes, SNPs in PURs ranged from 0.06-7.83%, and indels from 0.05-7.03% (**Table S26**), while the Y chromosome PURs contained the highest proportion of SNPs (30.51%). Within coding sequences (CDS), 2.25% of SNPs were in PURs, whereas intergenic regions accounted for 62.96%.

The Neighbor-Joining (NJ) tree revealed two major clades separating Asian and European pig populations (**Fig. 4D and Fig. S8**). The Asian clade primarily comprised Asian domestic pigs (AD) and AW, while the European clade included European domestic pigs (ED) and EW. Newer pig (NEW) breeds formed after the turn of the century, including Beijing Black and LW×Min (F1), are either hybrids or have substantial European breed introgression (*14*) (**Fig. 4E)**. For example, Shanghai White, Suhuai, and Yunong Black exhibited greater genetic similarity to European pigs, reflecting substantial European introgression. Principal component analysis (PCA) and Admixture analyses supported that NEW populations mainly occupy intermediate positions between Asian and European pigs (**Fig. 4F**). AD and AW exhibited higher genetic diversity (π=3.52×10^−3^ and 3.54×10^−3^, respectively), whereas EW showed the lowest diversity (π=1.65×10^−3^). The greatest genetic differentiation was observed between AW and EW (Fixation index (Fst) = 0.3679), whereas differentiation between AD and AW was much lower (Fst = 0.0445) (**Fig. 4G**), likely because ED and AD were independently domesticated from two wild boar populations that diverged approximately 1.23 Mya. Analysis of linkage disequilibrium (LD) decay and runs of homozygosity (ROH) showed that AD have lower LD and shorter ROH segments than ED, reflecting higher genetic diversity in AD populations (**Fig. 4H**). This difference in diversity likely results from intense selection in European breeds over the past several hundred years. The median fraction of runs of homozygosity (FROH) in AD was 0.206, compared with 0.415 in ED. NEW populations exhibited intermediate LD and a median FROH of 0.183 (**Fig. 4I**), consistent with their origin through crossbreeding between European and Asian pigs aimed at combining desirable economic traits from both lineages.

### Selective sweep analysis reveals key variants under selection in Asian pigs

During domestication, pigs acquired genomic and phenotypic traits distinct for Asian and European pigs (*15, 16*). We performed Fst analysis to identify genomic regions under selection between AD and ED, as well as EW and AW. This approach allows us to pinpoint candidate genes that may have been shaped by domestication, breeding practices, or adaptation to distinct environments between Europe and Asia. The average Fst values for AD versus ED and AW versus EW are 0.24 and 0.37, respectively, with the highest values also being 0.85 and 0.90, respectively (**Fig. 5A and Table S27**). Our analysis identified 472 genes showing signatures of selection in AD versus ED, and 585 genes in AW versus EW. Among the genes under selection, we located 3 previously unplaced (*SNURF, SNRPN, EHMT1*) and 5 (*CCDC194, LRRC69, SRRM5, SMR3A, ASDURF*) newly annotated genes. *MC1R* (chr6: 1,203,745-1,210,044bp) is the highest signal selected gene between ED and AD, known to regulate coat color (*17*); The second strongest signal is observed in *TBX19* showing strong differentiation between AD and ED. Interestingly, one missense variant in particular (**Fig. S9 A-B and Table S28**) may be associated with development, growth, and timidity traits in Asian pigs. The 4:g.87,337,427 C>T occurs in 24 ED and 968 AD, resulting in an arginine-to-histidine substitution. This mutation affects *TBX19*, a transcription factor crucial for pituitary corticotroph development and ACTH production, which regulates stress response and metabolic processes. Variants in *TBX19* may also influence behavior, including timidity and reactivity in AD, by modulating hormonal pathways linked to stress and temperament (*18*) (**Fig. S9C**). Predicted protein model showed a substantial deviation from the reference structure (RMSD: 17.504 Å), suggesting this mutation may impact on the protein structure and function (**Fig. S9D**); We co-located 249 selected regions (12.5Mb) in AD versus ED and AW versus EW groups, which most likely reflect ancestral divergence and natural selection before domestication. *PDPK1* is strongly differentiated in both AD versus ED and AW versus EW comparisons, suggesting a potential role in regulating growth and body size (*19*) (**Fig. S9E**). Genes such as *EHMT1, PDPK1, LINGO2* showed among the highest differentiation in Asian and European pigs. *EHMT1* shows strong signals of selection in both AD versus ED (Fst = 0.61) and AW versus EW (Fst = 0.80). This gene is a key regulator of brown adipose tissue cell fate and thermogenesis through the *PRDM16* complex, which likely explains its selective advantage in Asian pigs, known for higher fat deposition (*20, 21*). We detected 9 SNPs in CDS region (minor allele frequency, MAF>0.05), which showed strong linkage disequilibrium (**Fig. 5B**). A missense mutation (1:g.289,666,349 C>A) was identified predominantly in AD, resulting in an alanine-to-glutamic acid substitution (**Fig. 5, C and D**). Structural modeling predicts a substantial deviation from the reference protein (RMSD: 28.214 Å) (**Fig. 5E**), suggesting that this variant may alter *EHMT1* function, potentially affecting energy homeostasis, adiposity, and reproduction (*20, 21*).

**Fig. 5.**
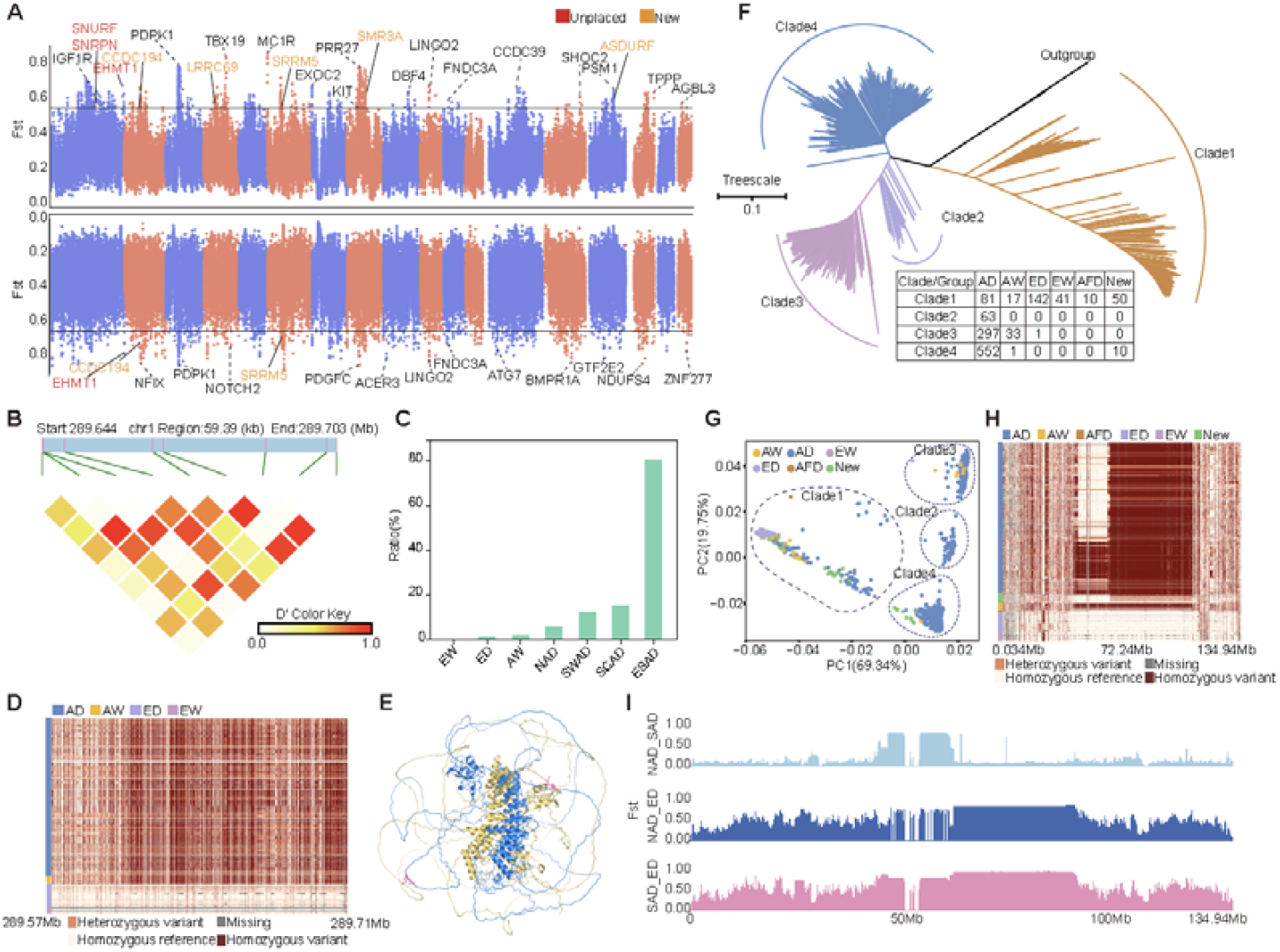
Selection signature analysis. **A**. Population divergence analysis using Fixation index (Fst) values calculated from genome-wide single nucleotide polymorphisms (SNPs), with the top 1% threshold indicated by the black horizontal line. Upper panel: Asian domestic pigs (AD) versus European domestic pigs (ED); lower panel: Asian wild boars (AW) versus European wild boars (EW). Orange indicates newly annotated protein-coding genes in T2T-Sscrofa; red indicates protein-coding genes from unplaced sequences in *Sscrofa*11.1 that were successfully positioned in T2T-Sscrofa. **B**. Linkage disequilibrium (LD) heatmap of SNPs within the coding sequence (CDS) of the *EHMT1* gene. **C**. Missense mutation frequencies across different pig populations. Population abbreviations: NAD, North AD; SWAD, Southwestern AD; SCAD, South-central AD; ESAD, Southeastern AD. **D**. Haplotype structure analysis of the *EHMT1* genomic region (chr1: 289.57-289.71 Mb). **E**. Predicted protein structural changes resulting from *EHMT1* point mutations. Yellow indicates wild-type (non-mutated) *EHMT1*; blue indicates mutated *EHMT1*. **F**. Neighbor-joining phylogenetic tree based on X chromosome SNPs. Population abbreviations: NEW: Newer pig populations formed after the turn of the century; AFD, African domestic pig; OG, outgroup. **G**. Principal Component Analysis (PCA) of domestic pig populations using X chromosome SNPs. **H**. X chromosome haplotype analysis. Population groupings as defined above. **I**. Fst-based population differentiation analysis of X chromosome variation among NAD, South AD (SAD), and ED populations.

### Divergent X chromosome haplotypes in Eurasian pigs

The X chromosome displayed a distinct phylogenetic structure (4 clades) that differed markedly from the autosomes (**Fig. S10**). While ED individuals showed similar distributions in autosomal and chrX clade1 (143 vs 142) (**Fig. 5F and Fig. S11**), AD representation was reduced on the X: 211 AD were assigned to autosomal clade1 (45.4%), while only 81 AD clustered in chrX clade1 (23.8%). Notably, 17 AW were uniquely found in chrX clade1, a pattern absent from the autosomes. Clade2 mainly comprises AW, and clade3 and 4 corresponded to north and south AD, and clade1 included nearly all European pigs, with some Asian samples (**Fig. 5G**).

Haplotype analysis revealed extensive X chromosome divergence among pig populations. South versus north AD carried distinct haplotypes in the 49.5-64.9 Mb region, while 64.9-96.2 Mb differentiated ED, EW, AD and AW (**Fig. 5H, Fig. S12 and Table S29**). Corresponding average Fst values were 0.15 (south AD vs. north AD), 0.49 (north AD vs. ED), and 0.57 (south AD vs. ED), respectively (**Fig. 5I**). Within 49.5-64.9 Mb, we identified 56 genes enriched for metabolic processes, cellular processes and reproduction (**Fig. S13 and Table S30**), harboring 198 missense mutations, and 50 predicted deleterious (SIFT<0.05) (**Table S31**).

These divergent haplotypes overlapped a large (~48 Mb) low-recombination block, with one haplotype prevalent in ED, EW and north AD, another in south AD and AW. The south Asian haplotype clustered with *Sus* species in Island Southeast Asia, consistent with introgression from an possibly extinct *Sus suid* lineage (*22*). Its current distribution likely reflects historical selection favoring northern latitudes, later transmitted to EW via Eurasian migrations. These results underscore the unique evolutionary dynamics of X chromosome haplotypes in Eurasian pigs.

## Discussion

Pigs are not only important domestic animals but also serve as valuable biomedical models due to their physiological and anatomical similarities to humans. The current reference genome (*Sscrofa*11.1) is extensively utilized across various research fields and serves as a fundamental resource for pig genomics studies (*4*). However, the pig reference genome (*Sscrofa*11.1) has remained largely unchanged for nearly eight years, which limits the resolution of genomic studies, particularly in complex or repetitive regions, due to incomplete assembly and remaining gaps. Although near T2T pig assemblies have been recently reported (*5, 23*), a fully complete T2T pig genome was still lacking with these genomes still containing incomplete centromere assemblies, unplaced sequences and gaps. Given the independent domestication histories, selective breeding, and phenotypic diversity among pig breeds (*24*), producing a high-quality, comprehensive reference genome remains a priority for both agricultural and biomedical research (*25*).

Advances in high-throughput sequencing technologies, including PacBio HiFi and ONT ultra-long, have made it feasible to assemble gapless, chromosome-scale genomes with high accuracy (*6*). Leveraging these technologies, we generated a complete T2T reference genome for the TJ Tabasco individual, including all 18 autosomes, the X chromosome, and the mitochondrial genome, supplemented with a public Y chromosome. This assembly (T2T-Sscrofa) captures PURs, complex repeats, and centromeric sequences that were missing from prior references, providing a more complete representation of the porcine genome.

A striking feature of T2T-Sscrofa is the remarkable expansion of acrocentric centromeres, up to 31.6Mb compared to the 2.5-6.5Mb centromeres of metacentric/submetacentric chromosomes. This expansion is driven by the accumulation of specific satellite repeats (pigcent4-6), resulting in regions with high repetitive content, weak Hi-C signals, and no detectable genes. Such centromere elongation, also observed in other mammals (*26*), likely arises from repeat amplification through unequal crossover or replication slippage and may influence chromosome stability and evolution (*7, 8*), particularly in lineages prone to Robertsonian fusions like the *Suids (15, 27*).

Using T2T-Sscrofa, we characterized SVs across Eurasian pigs, revealing extensive insertions, deletions, inversions, and duplications, many of which are enriched in PURs. These findings highlight the functional importance of repetitive and complex genomic regions, which harbor variants potentially affecting gene regulation, metabolism, and other phenotypic traits. In addition, the genome allowed us to resolve the unique Robertsonian fusion between chromosomes 15 and 17 in western EW (*28*). The assembly of a western EW with 2n = 36 chromosomes represents the first chromosome-level genome for this population and provides a foundation to study karyotype evolution, fertility in hybrids, and the functional consequences of chromosome rearrangements in *Suidae* species.

Population genomics analyses revealed substantial divergence between ED and AD, consistent with independent domestication events from distinct wild boar populations that diverged roughly 1 million years ago. AD display higher genetic diversity, lower linkage disequilibrium, and shorter runs of homozygosity compared to ED, likely reflecting the stronger intense artificial selection imposed on European breeds compared to Asian breeds over the past several centuries (*29*). Selective sweep analyses identified key genes under domestication and breed-specific selection, including *TBX19, EHMT1*, and *PDPK1*, affecting traits such as growth, fat deposition, reproduction, and behavior (*18, 19*). *EHMT1*, in particular, regulates brown adipose tissue cell fate and thermogenesis through the *PRDM16* complex, likely explaining its selective advantage in AD with higher fat deposition (*20*). A missense variant in *TBX19* is likely involved in more timid behavior (less activate and aggressive) in asian pig breeds (*30*). Interestingly, the South Asian X-chromosome haplotype clusters closely with haplotypes found in other *Sus* species from Island Southeast Asia, suggesting it may have originated from an ancient admixture event between South Asian wild boar and another, now extinct, suid lineage. A plausible explanation for its present-day distribution is that the haplotype conferred an adaptive advantage in northern latitudes, where it swept to fixation, and was later introduced into European wild boar populations through subsequent migrations across Eurasia.

Overall, the T2T-Sscrofa genome sequence provides an enhanced fundamental resource for the porcine genomics community, enabling pan-genome analyses, fine-mapping of structural and functional variants, investigation of karyotype evolution, single-cell transcriptomic studies, and precision genome editing. The inclusion of diverse wild boar and global domestic pig genomes further enhances its utility for understanding the genetic basis of economically and biologically important traits. This study underscores the value of complete, high-quality reference genomes in uncovering both the evolutionary history and the functional architecture of domestic animals.

